# Genetic and environmental sources of behavioral individuality: a test of the standard model

**DOI:** 10.1101/2025.08.27.672710

**Authors:** Zachary J. Sweep, Raphaël Royauté, Ned A. Dochtermann

## Abstract

Behavioral variation is typically assumed to arise from the combination and interaction of genetic and environmental variation. However, recent work with genetically identical individuals has found that substantial behavioral individuality is expressed even when genetic and environmental variation are negligible. This surprising result requires direct testing of our standard model for the sources of behavioral individuality. Here, we tested the standard model by comparing among-individual variation in highly inbred crickets versus outbred crickets. Comparing inbred and outbred lines allows for direct testing of the standard model by contrasting the magnitude of among-individual variation in a uniform versus varied genetic background. We found substantial and significant differences in among-individual variances, with among-individual variances being roughly three times greater in outbred versus inbred crickets (posterior probability, p = 0.974). Repeatability was also significantly different between inbred and outbred crickets (0.15 versus 0.41, respectively; p = 0.984). This result supports our standard model and suggests that the surprising expression of behavioral variation in clonal and parthenogenic species may represent an important but unique pathway for the expression of behavioral individuality.

## INTRODUCTION

Understanding how variation in behaviors is distributed among individuals has been a central topic of behavioral research over the last 20 years [1-4]. Among-individual variation has been identified under a variety of labels, including animal personality [e.g. 5], behavioral types [e.g. 6], and behavioral individuality [e.g. 7]. Regardless of the name used, behavioral individuality means that individuals differ from each other on average, even when each individual does not always show the exact same behavioral response from one time to the next [8]. A major finding of research into behavioral individuality is that among-individual variation is ubiquitous and accounts for ∼40% of the variation observed within populations [1]. Both genetic and environmental sources with long-term impacts have also been identified as being major contributors to patterns of behavioral individuality [reviewed in 9]. However, recent research has suggested that our standard model of how behavioral individuality arises may be incorrect or, at least, incomplete.

Specifically, under our standard model, phenotypic variation in behavior has been argued as emerging primarily from the combination and interaction of genetic and environmental variation [9, 10]. According to this model, behavioral individuality represents the sum and interaction of differential genetic contributions and long-term environmental effects. In this context, the relative magnitude of behavioral individuality is estimated based on variance components as the adjusted repeatability [1, 11-13]. Repeatability is the ratio of among-individual variation (V_I_) to total variation (V_T_), with among-individual variation being the genetic variance (V_G_) plus the variance due to developmental plasticity and what are known as permanent environmental effects (summed as V_PE_; [9, 14]):

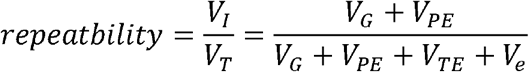

V_T_ includes among-individual variation, the effects of within-individual plasticity in response to unmeasured temporary environmental variation (V_TE_), and measurement and estimation error (V_e_). Repeatability connects to our classic model of behavioral individuality as emerging from genetic and environmental variation because both are represented in the ratio’s numerator.

This model of behavioral individuality is potentially incomplete because phenotypic variation has been identified in multiple systems where both genetic and environmental variation are assumed to be approaching zero. This “intra-genotypic variation” (IGV) has been identified in fruit flies [15-17], rodents [18], crustaceans [19], poecilid fish [20], and many other taxa. IGV has been argued to potentially be adaptive and representative of bet-hedging [21, 22], is partially inducible [15, 16], and there is genetic variation for the magnitude of IGV [16, 17]. The presence of IGV challenges our standard model if IGV manifests as among-individual variation and thus behavioral individuality when environmental variation is absent.

Work with Amazonian mollies (*Poecilia formosa*)—a species that reproduces parthenogenically—has found exactly this: IGV generates among-individual variation in the absence of apparent environmental variation. Moreover, the relative magnitude of this IGV is high enough that it suggests our classic model about the sources of among-individual variation is incorrect [7]. In a clonal species, like Amazonian mollies, if individuals of the same genotype are reared under standardized laboratory conditions, the variance components V_G_ and V_PE_ should both approach zero. While V_G_ will obviously approach zero in parthenogenic species, V_PE_ should do so under standardized laboratory conditions as well because its contribution to phenotypic variation is based on the amount of environmental variation present [23]. The amount of environmental variation present should, by definition, be small and approaching zero under standardized conditions. As these two variance components approach zero, repeatability, the relative magnitude of behavioral individuality, should also approach zero.

Surprisingly, repeatability of activity behaviors in Amazonian mollies does not approach zero. Instead, repeatability was initially estimated as ∼0.3 – 0.35 [7]. This is statistically indistinguishable from the meta-analytic mean of behavioral repeatability of 0.37, as reported by Bell, Hankson [1]. Subsequent work with mollies estimated repeatability of activity as ranging from ∼0.2 to 0.7 under a variety of contexts [24-26]. For repeatability to equal, or even exceed, the average for behaviors introduces the possibility that our standard model of the sources of phenotypic variation might generally be incomplete.

In contrast to the standard model, IGV is argued to arise due to stochastic variation in development [27] and developmental responses to unmeasurable micro-environmental variation [28]. The initial work by Bierbach, Laskowski [7] suggests that these mechanisms are sufficient to generate a similar relative magnitude of behavioral variation as proposed under the standard model. Alternatively, the evolution of mollies or parthenogenesis more generally might have included the acquisition of unique mechanisms for the generation of among-individual variation. Distinguishing between these explanations requires testing the classic model of behavioral variation in a wider range of systems, including non-parthenogenically reproducing systems.

Here we tested whether our standard model is sufficient to explain patterns of behavioral individuality in the banded cricket (*Gryllodes sigillatus*). To do so, we estimated the magnitude of among-individual variation in inbred and outbred lines of *G. sigillatus*. Importantly, because the standard model of behavioral individuality makes assumptions about the variance components summarized by repeatability, these variance components themselves must be estimated and compared.

## METHODS

To test predictions of the standard model, we measured the behavior of individuals from five inbred lines and one outbred line of *G. sigillatus*. Inbred *G. sigillatus* were the product of 23 generations of full-sibling matings starting in 2001 [29]. This resulted in an inbreeding coefficient of ∼0.99. Since then, for ∼50 generations, mating was random within lines. This random mating is expected to have led to an increased inbreeding coefficient due to drift and the stochastic loss of genetic variants. Consequently, individuals within lines can be considered effectively genetically identical. The outbred line was derived from individuals collected from Riverside, California in 2014 [30]. The outbred line was subsequently maintained with large population sizes and random mating. While this captive rearing necessarily resulted in some inbreeding, this line is nonetheless expected to have maintained greater genetic variation than is present in the inbred lines. Under the classic model of behavioral variation, we would therefore expect among-individual variation to be approaching zero in the inbred lines while being much higher in the outbred line.

### Behavioral Trials

We compared variation in exploratory propensity among and within lines. We measured exploratory propensity as an individual’s activity levels in an open field arena. All trials were conducted in an acrylic arena (60cm × 60cm and 15cm high) with a Plexiglas lid. The arena was split into four 30cm × 30cm arenas separated by a divider, allowing up to four crickets to be tested at one time. We excluded trials in which an individual moved less than 5 cm as in prior studies this would have been classified as a lack of exploration [31-34]. This resulted in the exclusion of 67 observations from our eventual analysis.

Individual crickets were left to rest for 30s under a 5-cm-diameter cup after being introduced into the lower right section of the arena (Figure S1). After 30s, we removed the cup and allowed the individuals to move freely through the arena for 220s. We measured each individual’s exploratory propensity by estimating the distance traveled (cm) by the cricket using Ethovision X (Noldus Information Technology). This behavioral protocol has previously been used with *G. sigilattus* and multiple other species of crickets [e.g. 32, 33, 35, 36].

Mass at the time of behavioral trials was recorded to the nearest mg and temperature at the time of testing measured to one-tenth of a degree Celsius. After each behavioral assay, arenas were thoroughly cleaned with 70% ethanol wipes to avoid accumulation of any chemical traces of conspecifics. All individuals were measured for a maximum of three repetitions, with some individuals measured fewer times due to escape or natural mortality (Table 1). Behavioral testing was conducted between June 2018 and May 2019 but, for logistical reasons (e.g. hatching and maturation times), testing was sequential across lines. In total, we included 611 behavioral trials across a total of 238 individuals in our analyses (Table 1).

**Table 1.** Number of individuals measured once or more from each inbred line and from the outbred line.

| Line | Individuals × number of repetitions |  |  | Total Trials |
| --- | --- | --- | --- | --- |
|  | ≥1 | ≥2 | =3 |  |
| A | 40 | 38 | 26 | 104 |
| E | 34 | 26 | 18 | 78 |
| F | 40 | 36 | 23 | 99 |
| G | 40 | 31 | 24 | 95 |
| H | 34 | 30 | 24 | 88 |
| Outbred | 50 | 50 | 47 | 147 |
| Total Individuals | 238 | 211 | 162 | Total: 611 |

### Statistical Analyses

Per the standard model for behavioral individuality, we predicted that the among-individual variance would be effectively zero in inbred but not in outbred crickets. Relatedly, we predicted that, if not zero, among-individual variances in the inbred lines would be substantially lower than for outbred crickets. This stems from the possibility that even if environmental variation is sufficient to generate behavioral individuality in the inbred crickets, the absence of genetic variation would still result in a lower among-individual variance. We were also interested in whether there were differences in behavioral averages and within-individual variances among lines but did not have *a priori* predictions about either. We also estimated adjusted repeatabilities for inbred and outbred crickets for comparison to other studies and, to better understand the magnitude of the variances, we estimated coefficients of variation (as percentages).

To determine whether among-individual variances differed between inbred and outbred crickets we fit a mixed-effects model using the MCMCglmm package in R [37]. With this model we allowed among- and within-individual variances to be estimated separately for inbred and outbred crickets. Individual cricket identity was fit as a random factor and temperature during testing and mass were included as centered fixed effects. Following Royauté and Dochtermann [38], we evaluated the posterior distribution of the difference between among-individual variances between the outbred and inbred crickets. Within-individual variances were also allowed to differ between the inbred and outbred crickets. Variances were determined to be significantly different if >95% of posterior estimate differences were greater for one group of crickets versus the other (i.e. posterior probabilities). We then ran additional models where means and among- and within-individual variances were estimated separately by line. MCMC analyses were run for 1.3 × 10^7^ iterations with a 3 × 10^6^ burn-in and a 10,000 iteration sampling interval.

Group means, variances, repeatabilities, and coefficients of variation are reported as the medians and 95% credibility intervals (CI) of posterior distributions [39]. Because of their magnitude, variances are reported as variance/10000 to aid interpretability.

There are two main caveats to our research here. First, our analyses here represent exploratory tests of the standard model. The data were originally collected to address other questions and were opportunistically used for our questions here. Second, because of their different origins, the founding population of the inbred lines likely had a different starting magnitude of genetic variation. While this may have been higher or lower than that of the outbred population, the high level of inbreeding would nonetheless be expected to drive variances in the inbred lines to zero while the random mating of the outbred line is expected to maintain more variation.

## RESULTS

The estimated median of among-individual variance in open field distance for outbred crickets was 2.3 (95% CI: 1.1:4.3) versus 0.81 (0:1.6) for the inbred crickets. As predicted by the standard model, the among-individual variance was significantly greater in outbred crickets (Figure 1A; outbred versus inbred median difference = 1.6 (0:3.6), 97.4% of posterior estimates were positive). Across posterior estimates, the among-individual variance of inbred crickets was ∼1/3 that of the outbred crickets (0.34, 0:1.02).

**Figure 1.**
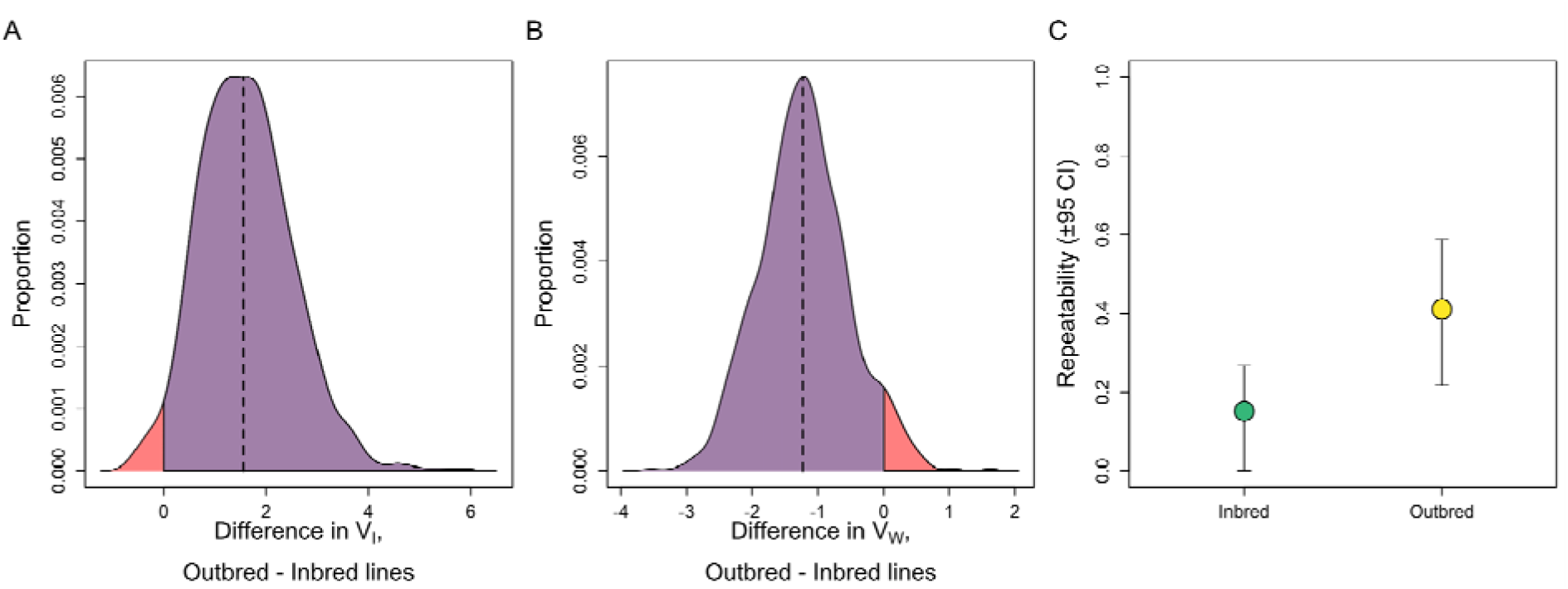
Relationships of among-individual variances, within-individual variances, and mean activity across inbred and outbred lines. (A) The MCMC posterior distribution of differences in among-individual variances (V_I_) between outbred and inbred crickets. The red portion of the distribution covers those estimates below zero. Among-individual variance was estimated as being greater in the outbred crickets in 97% of posterior estimates. (B) Posterior distribution of differences in within-individual variances (V_W_) between outbred and inbred crickets. The red portion of the distribution covers those estimates above zero. Within-individual variance was estimated as being smaller in the outbred crickets in 95% of posterior estimates. (C) Repeatabilities of activity in inbred (left) and outbred (right) crickets.

In contrast, the estimated within-individual variance was higher for inbred than outbred crickets. For outbred crickets, this variance was estimated as 3.4 (95% CI: 2.5:4.5) versus 4.6 (3.8:5.6) for the inbred crickets. The median difference estimated between inbred versus outbred crickets was -1.2 (-2.4:0.18), with within-individual variance for inbred crickets being greater in 95.2% of posterior estimates, a significant difference between the groups (Figure 1B). Across the posterior, the within-individual variance of outbred crickets was ∼3/4s that of the inbred crickets (0.73, 0.53:1.05).

Combined, the repeatability of open field distance was greater in outbred crickets than inbred crickets (median difference: 0.27, 0.03:0.48). This difference was significant (98.4% of posterior estimates) and repeatability of the outbred crickets was 0.41 (0.22:0.59) versus 0.15 (0:0.27) for the inbred crickets (Figure 1C).

Across lines, median distance traveled was highest in the outbred crickets (581.60cm, 487.10:664.18; Figure 2A), though one line of inbred crickets showed a similar degree of activity (line E: 519.95cm, 369.93:649.22). Across lines, two of the inbred lines (E and H) had among-individual variances that were roughly half of the estimated variance of the outbred crickets (Figure 2B). The inbred lines A, F, and G had among-individual variances that were approaching zero (Figure 2B). Within-individual variances differed across inbred lines but were generally of similar magnitude (Figure 2C). Among-individual coefficients of variation were around 1% for three of the inbred lines but, as was the case for among-individual variances, higher in lines E and H (Figure 3). Because of their relatively high among-individual variances—for the inbred lines—the coefficients of variation for these lines were similar to that of the outbred line, though both had uncertainties that extended to zero.

**Figure 2.**
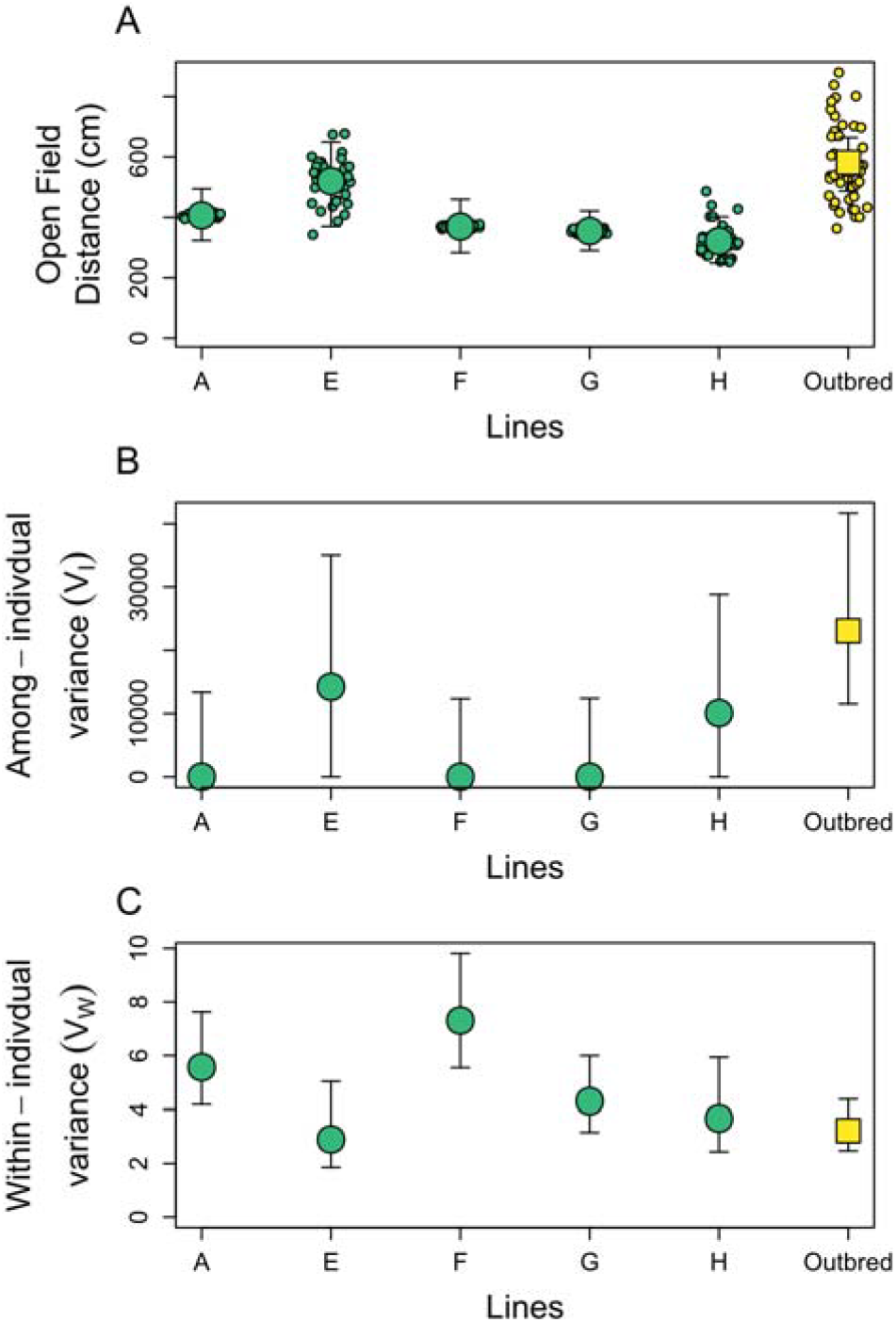
(A) Posterior median (large dots) and 95% credibility intervals (bars) of average activity—as measured by distance traveled in an open field (cm)—for the five inbred (green) and one outbred line of crickets (yellow). Small individual points are estimated values for individual crickets (best-linear unbiased predictors). (B & C) Median estimates (large dots) and 95% credibility intervals of among- and within-individual variances for the five inbred (green) and one outbred line (yellow) of crickets.

**Figure 3.**
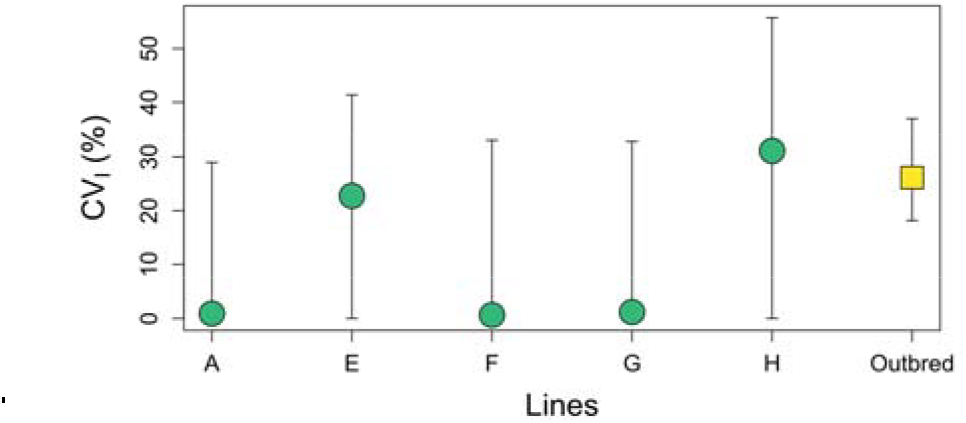
Posterior median estimates of coefficients of among-individual variation by line (as percentages) and 95% credibility intervals.

## DISCUSSION

Here, we found that the among-individual variation in outbred crickets was significantly greater than in inbred crickets (Figure 1A). Likewise, repeatability in the inbred crickets was substantially and significantly lower than in the outbred crickets (Figure 1C). These results support the standard model of behavioral individuality. This represents an important direct test of this model given recent observations that neither genetic nor environmental variation are necessary for behavioral individuality. While these sources of variation may not be necessary in parthenogenically reproducing systems, they were necessary here.

Interestingly, in two of our inbred lines (lines E and H), the estimated among-individual variance did not approach zero. Instead, for these lines, among-individual variances were estimated as around half the magnitude estimated in the outbred group (Figure 2B), as also reflected in the coefficients of variation (Figure 3). This may be the case for multiple non-exclusive reasons. First, if individuals from other lines were unknowingly introduced to these lines, inbreeding coefficients would be lower than expected, allowing for greater than expected among-individual variation. Second, since behavioral testing necessarily occurred in a disjunct manner across lines due to patterns of maturation not in our control, microenvironmental effects may have differed by line and resulted in greater than expected among-individual variation within only some lines. Third, if there is genetic variation in IGV, we would expect some lines to show greater among-individual variation. We cannot conclusively distinguish among these explanations at this time.

Consistent with microenvironmental effects explaining the higher among-individual variation in two lines, humidity in both the cricket housing and behavioral assay rooms is typically lowest during the winter (due to increased forced air heating). Line E, one of the high among-individual variance inbred lines (Figure 1D), was the only line measured during the winter. However, this explanation is not sufficient to explain the high among-individual variance estimated for line H, which was measured at the same time as line G. Moreover, given standardized housing, testing procedures, and temperature controls, we would expect micro-environmental effects to be random with regards to lines and so not sufficient to explain our results. Alternatively, lines E and H may produce greater amounts of IGV, suggesting genetic variation in IGV, as has been observed in other species [e.g. 17]. Unfortunately, while both these explanations are biologically interesting, the more parsimonious explanation is simply that genetic variation was unknowingly introduced into lines E and H at some point.

We also found significant differences in within-individual variances between the inbred and outbred crickets. We did not have *a priori* predictions regarding this variance component and so do not have a ready explanation for this finding. Unfortunately, relative to behavioral individuality, within-individual variation has been poorly studied despite its biological importance [40, 41]. Future research into within-individual variation may subsequently explain our findings here.

More generally, recent behavioral research has suggested that the standard model for sources of behavioral variation is insufficient [e.g. 7, 25, 26]. These were important findings and necessitate a reassessment and testing of our standard model. Our findings here indicate that the standard model may be sufficient, albeit not necessary. This conclusion suggests that findings elsewhere which supported a major role for IGV may instead be due to the specifics of an organism’s reproductive biology. Whether due to bet-hedging resulting in higher fitness [22], inter-group selection [as may be the case with mutation rates; e.g. 42], or other factors, mechanisms for the generation of variation may have been selected for in parthenogenically reproducing species. This is an intriguing possibility warranting further exploration. Similarly, Laskowski, Doran [43] argued that the unique biology of clonally reproducing vertebrates may reveal previously unrecognized biological insights. Regardless, the finding that our inbred crickets showed the expected and reduced among-individual variability supports the dominant assumptions of research regarding behavioral individuality.

## Data Accessibility

All data and code are available at https://osf.io/vk3a7/

## Author Contributions

ZJS and NAD conceptualized the project. NAD and RR jointly developed the analysis plan. RR led data collection. ZJS wrote the initial manuscript draft and all authors contributed to the final manuscript.

## Acknowledgements and Funding Statement

The authors thank Brady Klock, Ishan Josji, and Hannah Lambert for assistance with animal husbandry and behavioral trials. NAD, RR, and data collection were supported by NSF IOS-1557951. ZJS was supported by the NDSU Change RaMP Program and NSF DBI-2216605.

---

**Figure S1.**
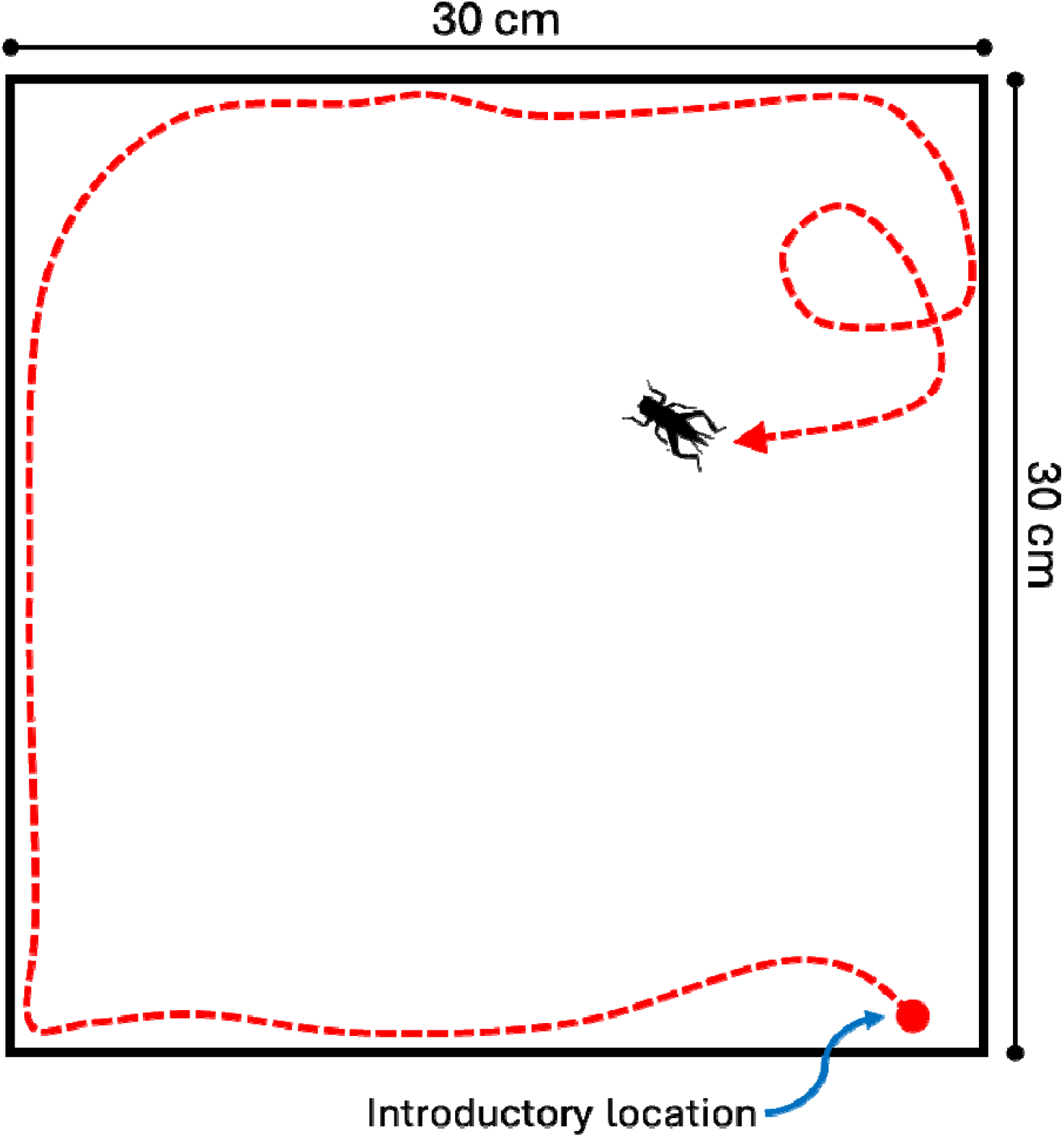
Schematic of the behavioral testing arena. The red arrow represents a possible path, the length of which (in cms) would represent an individual’s activity score.

